# Dual Effects of *Nanoviricides* Platform Technology Based NV-CoV-2 Biomimetic Polymer Against COVID-19

**DOI:** 10.1101/2021.11.24.469813

**Authors:** Anil Diwan, Ashok Chakraborty, Vijetha Chiniga, Vinod Arora, Preetam Holkar, Yogesh Thakur, Jay Tatake, Randall Barton, Neelam Holkar, Bethany Pond

**Affiliations:** Nanoviricides, Inc., Shelton, CT, USA; AllExcel, Inc., West Haven, CT, USA

**Keywords:** Remdesivir, COVID-19, SARS-CoV-2, Therapy, Vaccine, Antiviral, NV-CoV-2 (387-Polymer), Nanoviricides

## Abstract

Remdesivir (RDV) is the only antiviral drug so far approved for COVID-19 therapy by the FDA. However its efficacy is limited *in vivo* due to its low stability in presence of plasma. This paper compared the stability of RDV encapsulated with our platform technology based polymer NV-387 (NV-CoV-2), in presence of plasma *in vitro* and *in vivo*. Furthermore, a non-clinical pharmacology studies of NV-CoV-2 (Polymer) and NV-CoV-2-R (Polymer encapsulated *Remdesivir*) in both NL-63 infected and uninfected rats were done. In an *in vitro* cell culture model experiment, antiviral activity of NV-CoV-2 and NV-CoV-2-R are also compared with RDV.

The results are (i) NV-CoV-2 polymer encapsulation protects RDV from plasma-mediated catabolism *in vitro* and *in vivo*, too. (ii) Body weight measurements of the normal (uninfected) rats after administration of the test materials (NV-CoV-2, and NV-CoV-2-R) show no toxic effects on them. (iii) NL-63 infected rats body weights and their survival length were like uninfected rats after treatment with NV-CoV-2 and NV-CoV-2-R, and the efficacy as an antiviral regimen were found in the order as below: NV-CoV-2-R > NV-CoV-2 > RDV.

In brief, our platform technology based NV-387-encapsulated-RDV (NV-CoV-2-R) drug has a dual effect on coronaviruses. First, NV-CoV-2 itself as an antiviral regimen. Secondly, RDV is protected from plasma-mediated degradation in transit, rendering altogether the safest and an efficient regimen against COVID-19.

## Introduction

There are seven coronaviruses, so far, were identified, that infect humans, and only 4 of them belong to the beta family of coronavirus (HCoV-HKU1, SARS-CoV-2, MERS-CoV and SARS-CoV) [1]. SARS family were known to cause severe respiratory disease in humans. In fact, SARS-CoV-2 infection caused a pandemic COVID-19 disease with high morbidity and mortality [2]. *Nanoviricide* is a platform-technology-derived biomimetic polymer that can bind to a virus-specific ligands [3]. The flexible polymer backbone is comprised of a polyethylene glycol (PEG) and alkyl pendants. Different virus-specific ligands can be conjugated to this PEG-based polymer backbone to produce different individual drug substances that specifically target different viral pathogens.

Viruses bind to specific cell surface ligands in order to attach to cells and internalize. A *nanoviricide* is designed to act like a decoy of a human cell. When the virus sees the appropriate ligand displayed on a *nanoviricide* micelle, the virus is believed to bind to it. The *nanoviricide*, being flexible, could allow maximization of binding by spreading onto the virus particle, and potentially leading to fusion with the lipid-coated virus surface by a phase-inversion, wherein the fatty core of the *nanoviricide* could merge with the viral lipid coat and the hydrophilic shell of the *nanoviricide* could become the exterior of the particle, thus potentially engulfing the virus [4]. In the process, the coat proteins that the virus uses for binding to cells would be expected to become unavailable. This highly targeted attack may lead to loss of the coat proteins and the *nanoviricide* may further dismantle the engulfed virus capsid. The loss of virus particle integrity could render the virus non-infectious. This proposed mechanism of action is shown graphically in **Figure 1**.

**Figure 1:**
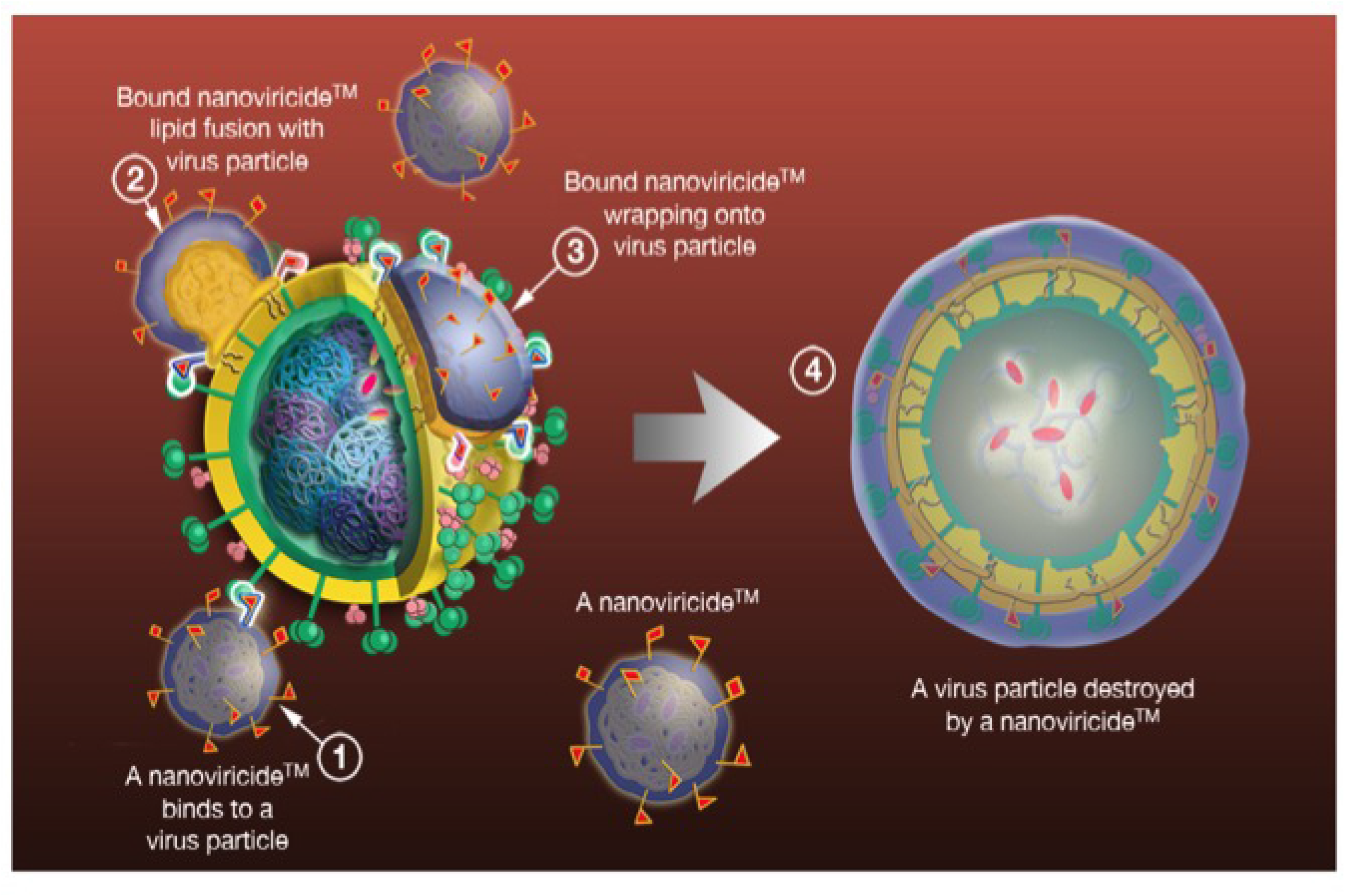
Graphic Representation of the mechanism of action of *nanoviricide* binding and inactivation of viruses

Support for this mechanism is shown in the electron photomicrographs in **Figure 2** in which murine cytomegalovirus (MCMV) was incubated with a *nanoviricide*. As can be seen in **Figs. 2B & C**, the binding of the *nanoviricide* to the CMV results in loss of the viral envelope; the resulting CMV naked capsids are non-infectious. Thus, *nanoviricide* binding renders the CMV not only inactive but appears to result in the disruption of capsid organization [5]. Given this proposed mechanism of action, *nanoviricides* are not expected to interfere with the intracellular replication of the virus to any appreciable extent.

**Figure 2:**
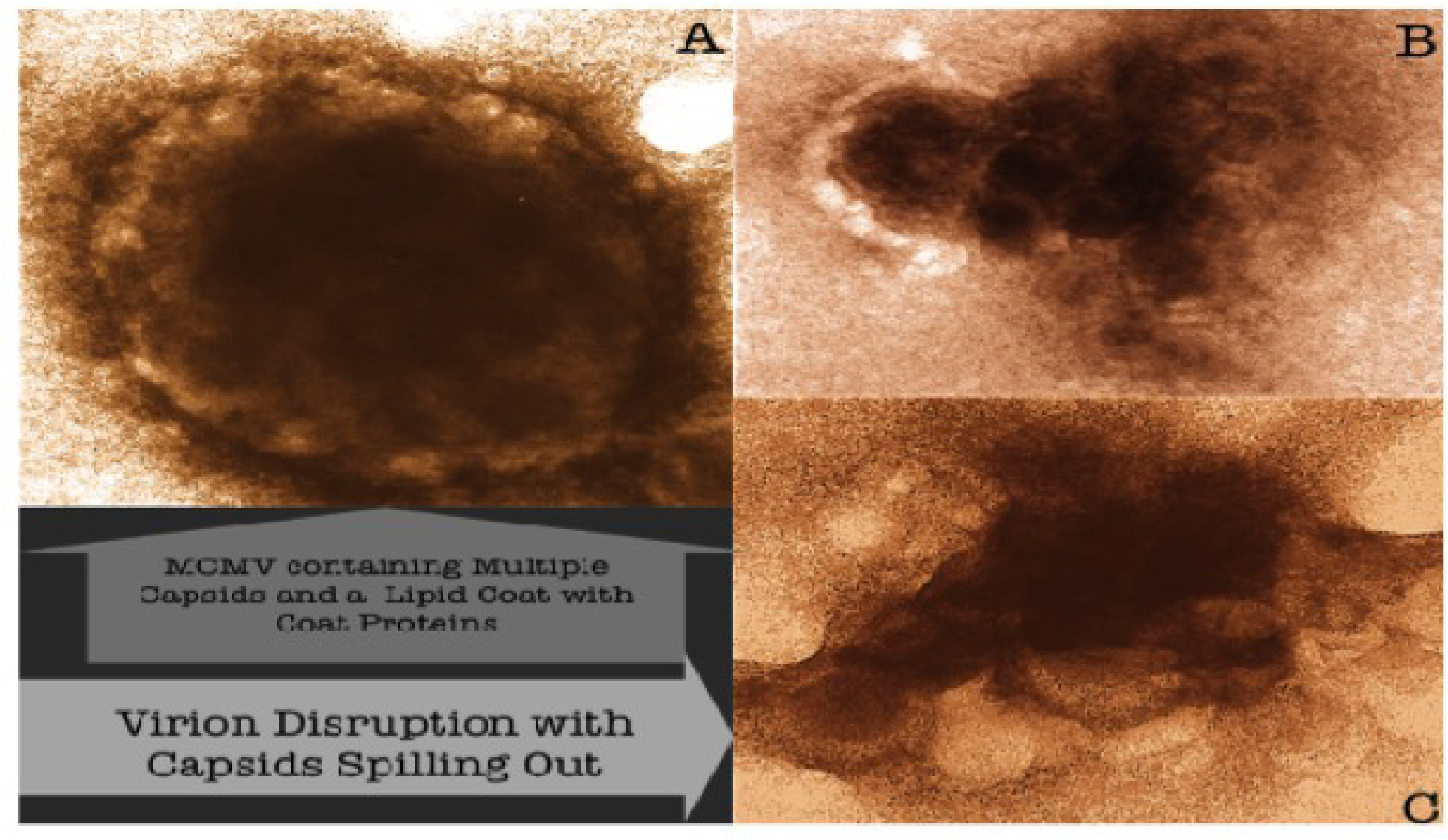
Effects of two different *nanoviricides* binding to murine cytomegalovirus (MCMV) A) Control treated Virion; B&C) MCMV virions treated with two different *nanoviricides*

Here we will be reporting the various aspects of this drug: (i) It’s antiviral activity *in vitro* cell culture model and *in vivo* rat animal model; (ii) It’s capability of increasing the half-life of RDV in presence of plasma *in vitro* and *in vivo;* (iii) and some safety aspects of this drug.

## Methods

### Materials

The drug substance, NV-CoV-2 was made in our lab. Based on polyethylene glycol of 22 repeat units (m) substituted by 2 hexadecylamine (R_1_) the theoretical molecular formula for the repeat unit of the polymeric drug substance (NV-CoV-2) is C_104_H_188_N_2_O_44_S_4_ *(Much details of the chemistry of the drug can not be disclosed at this point due to patent right obligation)*. It is an off-white, waxy non-crystalline semi-solid at room temperature, has no noticeable smell and it has a slight sour test.

Sources of other reagents, like RDV, GS-441524 and their internal standard (^13^C_6_- and ^13^C_5_-isotopes, respectively) were form *Medko Biosciences, Inc.* (USA), and *AlsaCHIM* (USA) respectively.

### Quantitative composition of Drug Product NV-CoV-2 Solution for Injection

The drug product NV-CoV-2 solution for injection is an injectable dosage form that will be supplied in a single vial as 10 ml of a 50 mg/ml solution of the drug substance and 40mg/ml D-mannitol (or quantity needed for osmolality) in water for injection. This solution is ready for injection to deliver 450 mg of drug substance NV-CoV-2 in 9 ml.

### Pharmacokinetics

Since NV-CoV-2 is composed of a flexible PEG-based polymer backbone, conventional methodologies to measure the pharmacokinetic properties of small molecules are not suitable for use with NV-CoV-2. We have developed a Direct ELISA to quantitatively detect NV-CoV-2 in biological matrices in order to measure its pharmacokinetic properties.

We have performed a non-GLP study to determine the levels of systemic exposure of NV-CoV-2 when NV-CoV-2 drug is administered to rats once daily for 5 days (0,1,3,5 and 7) over a 8-day period. Detection and Quantification of NV-CoV-2 in plasma samples was done by Direct ELISA. This assay method is a colorimetric ELISA to measure the levels of NV-CoV-2 in processed plasma samples. As described in the **Table 1,** thirty-six animals (three/sex/group) were administered NV-CoV-2, at three different doses each, or vehicle once daily, 5 times over an eight-day period on days 0,1,3,5 and 7. Injection on Day 0 is the 1st injection and on Day 7 is the 5th injection. Each compound was delivered via slow-push IV injection. Blood Samples for systemic exposure were collected at 0, 0.08, 0.5, 1, 2, 4, 8 and 24 hours after 1st and 5th injection of drug administration. Blood samples were taken from one animal per sex in each test article treated group at each time point. The same animal was used to draw blood across all the time points after Day 0 and Day 7 injections.

**Table 1:** Drug Injection Protocol.

| Group | Drug | Dose (mg/kg) | Dose Volume (ml/kg) | Concentration (mg/ml) NV-CoV-2 |
| --- | --- | --- | --- | --- |
| 1 | NV-CoV-2 High | 320 | 10 | 32 |
| 2 | NV-CoV-2 Medium | 160 | 10 | 16 |
| 3 | Vehicle | 0 | 10 | 0 |

Detection and Quantification of NV-CoV-2 in the plasma samples from each group was done by using a direct ELISA. The biotinylated detection antibody, 5G8-Biotin, used in this assay has been shown to be specific to NV-CoV-2. The Direct ELISA assay for NV-CoV-2 was previously characterized for linearity and detection limit. The limit of detection (LOD) of this assay is 0.25 μg/ml.

### *In vitro* antiviral activities of NV-CoV-2

The antiviral efficacy and potency, and the cytotoxicity of the antiviral compound NV-CoV-2 in cell cultures were studied. Like SARS CoV-2, CoV-NL63 binds to angiotensin-converting enzyme 2 (ACE2) for entry into cells [6]. Therefore, CoV-NL63 could be considered as a better surrogate virus for studying SARSCoV-2 *in vitro* and *in vivo* [7].

Rhesus Monkey Kidney Epithelial (LLC-MK2) cells and Human Fetal Lung Fibroblast (MRC-5) cells were obtained from ATCC and grown in Dulbecco’s Modified Eagle Medium (DMEM), supplemented with 10% fetal bovine serum (FBS), and 1% Penicillin-Streptomycin-Amphotericin B according to provider’s instructions. Human Coronavirus 229E (HCoV 229E) was purchased from ATCC and propagated on MRC-5 cells at 34°C for 5-7 days according to manufacturer’s instructions to produce virus stocks. These CoV 229E virus stocks were aliquoted and stored at -80°C until needed. Human Coronavirus NL63 (HCoV NL63) was purchased from ATCC and propagated on LLC-MK2 cells at 34°C for 6-9 days according to manufacturer’s instructions to produce virus stocks. These CoV NL63 virus stocks were aliquoted and stored at -80°C until needed.

In brief, LLC-MK2 cells (for CoV NL63 virus) and MRC-5 cells (for CoV 229E virus) were plated in 96-well black, clear bottom plates at a density of 30,000 cells per well in 100 μl media for 24hr prior to infection. Stock compounds were made at a concentration of 20 mg/ml in various vehicles. On the day of infection, compounds were serially diluted in DMEM medium 4x the required dose (i.e., a final concentration of 320 μg/ml was initially made at 1280 μg/ml). Virus was then diluted to the appropriate infection PFU/well dose: CoV NL63 and CoV 229E MOI=0.01. In a separate 96-well plate 60 μl of compound were added to 60 μl of virus and incubated for 1 hour at 34°C. After incubation, 100 μl/well of compound:virus mixture was added to each well of the cell plates, for a total of 200 μl and 1x dilution of compound. Plates were incubated at 34°C for appropriate amount of time for each virus.

We used the Promega CellTiter-Glo luminescent cell viability assay to test for compounds cytotoxicity. This assay allows the differentiation between death caused by compound toxicity and actual virus suppression/cytopathic effect. Media were removed, and cells were washed with PBS before adding 100 μl of serum-free media to each well. The assay was performed according to manufacturer’s instructions. The concentration of compound that causes reduction of cell viability by 50% is termed Effective dose or concentration 50 (EC50).

#### Virus cytopathic effect (CPE) ASSAY

A cell-based viral CPE assay in 96-well black, clear-bottom plates was developed which allows a higher-throughput screening of compounds. Virus (CoV229E and CoVNL63) were pre-incubated with compounds and then added to MRC-5 cells or LLC-MK2 cells, respectively, and incubated for 5-7 days. This gives the virus time to replicate and cause cell death (cytopathic effect). Relative CPE levels can be compared between all groups to identify any reduction in viral replication, as a reduction in CPE compared to untreated infected controls suggests a decrease in virus production/growth/spread.

### *In vivo* antiviral activities of NV-CoV-2: Dose-response of NV-CoV-2 in rats infected with CoV-NL63

Male and female Sprague Dawley rats, 8 to 9 weeks old, were infected with 10^4^ Cov- NL63 viral particles directly into the lungs. All untreated rats infected in this manner succumb to the disease in 5 to 6 days and, hence, used as a model for evaluation of the efficacy of NV-CoV-2. NV-CoV-2 and Vehicle were administered by tail vein I.V. injection (10 ml/kg), once per day, on Days 0, 1, 3, 5, and 7. RDV was administered by tail vein I.V. injection as per the standard administration protocol of twice for the first day and once daily after that for seven days. A group of infected, untreated rats was included as an additional control. The groups of animals and the treatments were shown in the **Table 2** below.

**Table 2:** Treatment Groups and Regimen:

| Treatment Group and Regimen | Concentration (mg/mL) | Dosage (mg/kg) | Total Dosage (mg/kg) | # of Rats |  |
| --- | --- | --- | --- | --- | --- |
|  |  |  |  | M | F |
| Untreated | 0 | 0 | 0 | 5 | 4 |
| Vehicle<br>i.v. Days 0,1,3,5,7 | 0 | 0 | 0 | 8 | 8 |
| Remdesivir<br>i.v. Days 0 (2x),<br>1,2,3,4,5,6,7 | 1.0 | 10 | 90 | 8 | 8 |
| NV-CoV-2 Med<br>i.v. Days 0,1,3,5,7 | 16 | 160 | 800 | 8 | 8 |
| NV-CoV-2 High<br>i.v. Days 0,1,3,5,7 | 32 | 320 | 1600 | 8 | 8 |

The rats were observed daily for clinical and behavior changes and body weight; moribund rats were sacrificed. The primary end points were survival and body weight loss.

### Effect of Number of Days of Dosing of NV-CoV-2 in Rats Infected with CoV-NL63

Male and female Sprague-Dawley rats, 8 to 9 weeks old, were infected with 2 x 10^4^ Cov- NL63 viral particles directly into the lungs. All untreated rats infected in this manner succumb to the disease in 5 to 6 days and, hence, used as a model for evaluation of the efficacy of NV-CoV-2. NV-CoV-2 was administered by tail vein I.V. injection, once per day, for different numbers of days of treatment, 1, 3 or 5 days beginning on Day 0. The vehicle control treated rats were treated for 5 days on Days 0, 1, 3, 5,and 7. A group of infected, untreated rats was included as an additional control. The groups of animals and the treatments were shown in the **Table 3** below.

**Table 3:** The groups of animals and the treatment protocol.

| Group | NV-CoV-2<br>mg/ml | Day of<br>Injection | No of<br>injections | Injection<br>volume<br>ml/kg | NV-CoV-2<br>Dose per<br>Injection<br>mg/kg | Total<br>NV-CoV-2<br>injected<br>mg/kg | No of<br>Animals |  |
| --- | --- | --- | --- | --- | --- | --- | --- | --- |
|  |  |  |  |  |  |  | M | F |
| NV-CoV-2<br>1 day | 50 | 0 | 1 | 12 | 600 | 600 | 7 | 7 |
| NV-CoV-2<br>3 Days | 50 | 0, 1, 3 | 3 | 12 | 600 | 1,800 | 7 | 7 |
| NV-CoV-2<br>5 Days | 50 | 0, 1, 3, 5, 7 | 5 | 12 | 600 | 3,000 | 7 | 7 |
| Vehicle<br>Control |  | 0, 1, 3, 5, 7 | 5 | 12 | 0 | 0 | 4 | 4 |
| Infected<br>Untreated |  |  | 0 | 0 | 0 | 0 | 2 | 2 |

The infected rats were treated with the same dose per injection, 600 mg/kg of NV-CoV-2 for the NV-CoV-2 treatment groups. Only the number of treatment days varied. The rats were observed daily for clinical and behavior changes, body weight; moribund rats were sacrificed. The primary end points were survival and body weight loss.

### Protection Efficiency of NV-CoV-2 encapsulation of RDV (NV-CoV-2-R) from plasma *in vitro*

First, a standard curve of RDV was determined by using LC-MS at different concentrations of the standard solutions ranging from 0-5 ng/uL in DMSO:MeOH (1:9). As an internal standard (ISTD), ^13^C_6_ –RDV (2.5 ng/uL) was used. Extraction of RDV for LC-MS assay was done by using acetonitrile. Concentrations of RDV compared to their ratio with the ISTDs (^13^C_6_-RDV) was used to generate a standard curve for RDV.

Secondly, ten microliter of the test materials (20 ng/uL) was incubated with 30 uL rat plasma for different time points (as indicated in the Result section), and then extracted with acetonitrile (100 uL). The mixture was centrifuged for 10 mins at 10,000 x g to separate the supernatants from the precipitates. The supernatants containing either RDV was determined by LC-MS chromatography as described Elsewhere [9].

The ratio of RDV and its isotope, ^13^C_6_-ISTD (as an internal standard) was calculated. The amount of RDV was determined using the linear equation derived from their respective standard curve. The values were normalized using the dilution factor used for the original plasma sample.

### Protection Efficiency of NV-CoV-2 encapsulated RDV (NV-CoV-2-R) *in vivo*

Thirty-six Sprague Dawley male rats (Taconic Biosciences, USA) (three/ in control and in treatment groups) were administered with NV-CoV-2 or NV-CoV-2-R, once per day for 5 days (0, 1, 3, 5, and 7) over a 7-day time period. NV-376 (RDV-in-SBECD; Gilead) was given as two doses on day 1 followed by a daily doses through day 7. DMSO was used as a vehicle for the control group. Injection on day 0 is considered as a 1^st^ injection and on day 7 is the 5^th^ injection. Each compound was delivered via slow-push IV injection. Blood samples for systemic exposure assay were collected at 0, 0.08, 0.5, 1, 2, 4, 8 and 24 hours after 1^st^ and 5^th^ injection of the drugs.

Blood samples were taken from one animal in each test article treated group at each time point. Same animal was used to draw blood across all the time points after “day 0” and ”day 7” injections. The detailed procedures and study design were stated in elsewhere [9]. RDV values obtained in rat plasma (mg/mL) after 1^st.^ and 5^th^ injection of the drugs were normalized by dividing with the amount of RDV administered (mg/kg of rat body weight).

### Safety pharmacology studies

We have conducted safety pharmacology studies; core battery tests as defined in the ICH S7A guidelines on the respiratory and central nervous systems in the rat and the cardiovascular system in the monkey after intravenous administration of NV-COV-2.

#### Central Nervous System and Respiratory System

Evaluation of Pulmonary and Neurobehavioral Functions Following Intravenous Administration of NV-CoV-2 in Conscious Rats (Calvert Laboratories, Inc., Scott Township, PA) in 24 male rats (Strain/Substrain: Crl:CD^®^(SD)] of age range 6 weeks and with each body weight of 162-199 Each group contains 6 animals. Route of administration of the drug is i.v, for once and dose volume is 5 ml/kg **(Table 4).**

**Table 4:** Test Article Administration: Group Assignments and Dose Levels:

| Group | Treatment | Dose<br>(mg/kg) | Volume<br>(mL/kg) | Concentration<br>(mg/mL) | No. of Animals |
| --- | --- | --- | --- | --- | --- |
|  |  |  |  |  | Male |
| 1 | Vehicle | 0 | 5 | 0 | 6 |
| 2 | NV-CoV-2 Low | 25 | 5 | 5 | 6 |
| 3 | NV-CoV-2 Medium | 50 | 5 | 10 | 6 |
| 4 | NV-CoV-2 High | 100 | 5 | 20 | 6 |

### Pulmonary Evaluation

Rats were trained for two days in the head-out plethysmograph chamber, prior to the experiment. On the day of dosing, each animal was weighed and placed in the plethysmograph chamber (SOP PHA-EQP-96) and baseline respiratory parameters were obtained for 5 minutes following a stabilization period. The rat was then removed from the chamber and dosed as per group assignments. After dosing, each animal was returned to its designated plethysmograph chamber, allowed to stabilize, and the respiratory parameters were measured for 5-minute intervals at 15 min, 1, 2 and 4 hours following administration of test article/vehicle. Respiratory parameters were measured at ±5 minutes for the 1-hour evaluation and ±15 minutes for the 1, 2 and 4 hours post-dose evaluations. After each reading the animals was removed from the plethysmograph chamber and returned to its home cage. The following parameters were recorded (SOP PHA-SOF-90) using the Ponemah Physiology Platform (Ponemah v.5.20 Pulmonary): Respiratory rate; Tidal Volume, and Minute Volume.

### Neuropharmacology Evaluation

Modified Irwin/Functional Observations Battery test was used for the neuropharmacology evaluation. At approximately 15 minutes, 1 hour and 4 hours after dosing, and after the completion of the pulmonary evaluation, animals were placed (in groups) in a fixed environment consisting of a Plexiglas® enclosure, fitted with a lid. The enclosure was placed on absorbent paper that detects excretions. In this environment, the rats were free to move about.

Observances for the presence of the following symptoms were made as a group in the Plexiglas® enclosure. Animal number was documented when any of the following symptoms were noted except for excretion that were documented as a group observation: Seizures/convulsions (Awareness reaction); Body tremors (Motor activity); Ataxia (Piloerection); Abnormal posture (Stereotypy); Excretion (Decreased respiration).

Startle response was tested by clapping the hands immediately above the open square and noticing the response. Each animal was individually removed from the square and examined for vocalization, irritability, increased secretion of saliva and/or tears and for abdominal tone by feeling the abdominal muscles. Loss of righting reflex was examined by placing each animal on its back. Pupil size was examined individually for mydriasis or miosis under a magnifying glass. Increased or decreased nociceptive response was tested by tail pinch and corneal reflex and pinnal reflex was checked by a cotton swab and a blunt probe respectively. Immobility and grip strength for each animal was tested on an inclined wire screen.

**Rectal body temperatures** were taken at approximately 1 and 4 hours after dosing from all animals using a YSI Precision Digital Thermometer (Model 4600).

#### Cardiovascular System

Evaluation of Cardiovascular Function Following Administration of NV-CoV-2 in Conscious Telemetered Cynomolgus Monkeys (Calvert Laboratories, Inc., Scott Township, PA.) in the animal test system (Species: Monkey; Breed: Cynomolgus (*Macaca fascicularis*); Total Number /Sex): 4 Male; Age Range: 2-5 years; Body Weight Range: 5.1 to 6.0 kg prior to treatment). Test article administration and group assignments are shown in the **Table 5.** Dose administration are shown in **Table 6.**

**Table 5:** Test article administration.

| **Treatment** | **Dose Level (mg/kg)** | **Dose Volume (mL/kg)** | **Dose Conc. (mg/mL)** |
| --- | --- | --- | --- |
| Vehicle | 0 | 5 | 0 |
| NV-CoV-2 | 25 | 5 | 5 |
| NV-CoV-2 | 37.5 | 5 | 7.5 |
| NV-CoV-2 | 50 | 5 | 10 |

**Table 6:** Dose Administration Schedule (Latin Square Design)

| Treatment | Dose (mg/kg) | Day of Dosing<sup>a</sup> |  |  |  |
| --- | --- | --- | --- | --- | --- |
|  |  | Animal #1 | Animal #2 | Animal #3 | Animal #4 |
| Vehicle | 0 | Day 1 | Day 8 | Day 15 | Day 22 |
| NV-CoV-2 | 25 | Day 22 | Day 1 | Day 8 | Day 15 |
| NV-CoV-2 | 37.5 | Day 15 | Day 22 | Day 1 | Day 8 |
| NV-CoV-2 | 50 | Day 8 | Day 15 | Day 22 | Day 1 |

<sup>a</sup> A washout period of at least 3 days was allowed between each dose administration.

NV-CoV-2 formulations were infused intravenously for only one time using a calibrated infusion pump into a saphenous vein over a period of 30 (± 2) minutes using a percutaneous i.v. catheter. Animals were restrained during dosing. Animals were trained to the restrainers prior to dosing.

### Method of Cardiovascular Evaluation

Data were collected for approximately 24 hours before the first dosing day in order to establish acceptable baseline parameters for each animal. Baseline data were retained in the study file, but not included in the final report. Data were also collected for at least 24 hours prior to each subsequent dose to determine pre-dose values. Following dose administration, parameters were recorded continuously for approximately 24 hours. A washout period of at least 6 days was allowed between test article doses.

The following parameters were analyzed using the Ponemah 5.20 SP9 and ECG Analysis Module v. 5.30 software (SOP PHA SOF-106): Arterial Blood Pressure; Heart Rate (HR); ECG: Body Temperature. One-minute tracings of the ECGs at 15 minutes prior to dosing and at 5 and 30 minutes, 1, 2, 4, 8, 12 and 24 hours post-dose were printed out from the software acquisition and morphological changes were evaluated by a board-certified veterinary cardiologist.

### Immunogenicity

Since NV-CoV-2 is administered I.V., it is important to determine if there is any evidence of immunogenicity. A study was conducted for the detection of Anti-NV-CoV-2 antibodies in Rats with serum samples collected at 14 and 28 days following the start of 6 sequential I.V. injections of NV-CoV-2 over a period of 10 days, as described in the **Table 7** below.

**Table 7:** Immunogenicity study Design.

|  | Groups | NV-CoV-2<br>mg/ml | R<br>mg/ml | Days of<br>Injection | Volume<br>per kg | NV-CoV-2<br>Dose mg/kg | No of Animals |  |
| --- | --- | --- | --- | --- | --- | --- | --- | --- |
|  |  |  |  |  |  |  | Male | Female |
| 1 | NV-CoV-2 | 50 | 0 | 0, 1, 3,<br>5,7,9 | 12 | 600 | 4 | 4 |
| 2 | Vehicle | 0 | 0 | 0, 1, 3,<br>5,7,9 | 12 | 0 | 4 | 4 |
| 3 | Naive |  |  |  |  |  | 2 | 2 |

Detection and quantification of anti-NV-CoV-2 antibodies in serum samples was performed using an Indirect ELISA assay. This assay method is a colorimetric ELISA to measure the levels of anti-NV-CoV-2 antibodies in liquid samples. The antigen (NV-CoV-2) is first immobilized onto each well of a 96-well assay plate and then incubated with diluted subject serum samples or positive control, normal serum spiked with rat Anti-NV-CoV-2 antibodies. The bound Anti-NV-CoV-2 antibody is then detected by incubating with 1:1 mixture of HRP-conjugated Goat Anti-Rat IgG and IgM antibodies and quantitated with a chromogenic HRP substrate (TMB). A purified preparation of Monoclonal Antibodies, (IgG and IgM), host species-Rat, specific to NV-CoV-2 was used as positive control in naive subject matrix to determine the sensitivity. This Indirect ELISA method is specific and selective to the NV-CoV-2.

## Results

### 1. Pharmacokinetics

A typical time course of drug plasma concentration was observed over 24 hours. Results of the analyses are summarized in the **Table 8** below with Observed Maximum Plasma Concentration of NV-CoV-2 (C_max_), and the time to reach C_max_ (t_max_). The dose is expressed in mg/kg while the (C_max_) and (t_max_) are expressed in mg/ml and hours respectively.

**Table 8:** Results of NV-CoV-2 toxicokinetics analyses in groups treated with NV-CoV-2. Observed Maximum plasma concentration (C_max_) and time to reach C_max_, (t_max_). The C_max_ and t_max_ are expressed in mg/ml and hours respectively.

| | Dose<br>(mg/kg)<br>NV-CoV-2 | Dose<br>Volume<br>(ml/kg) | NV-CoV-2<br>Concentration<br>mg/ml | $t_{max}$<br>Hrs | $C_{max}$<br>mg/ml | +/- SD<br>(mg/ml) |
| --- | --- | --- | --- | --- | --- | --- |
| NV-CoV-2-High Male-Day1 | 320 | 10 | 32 | 8 | 0.64 | 0.06 |
| NV-CoV-2-High Male-Day 7 | 320 | 10 | 32 | 0.08 | 2.60 | 0.29 |
| NV-Cov-2-Medium Male-Day1 | 160 | 10 | 16 | 24 | 1.39 | 0.13 |
| NV-CoV-2-Medium Male-Day 7 | 160 | 10 | 16 | 4 | 1.55 | 0.07 |
| NV-CoV-2-High-Female Day1 | 320 | 10 | 32 | 4 | 2.48 | 0.10 |
| NV-CoV-2-High-Female Day 7 | 320 | 10 | 32 | 4 | 1.02 | 0.03 |
| NV-CoV-2-Medium-Female Day1 | 160 | 10 | 16 | 8 | 0.50 | 0.06 |
| NV-CoV-2-Medium-Female Day 7 | 160 | 10 | 16 | 4 | 0.08 | 0.05 |

Detection and Quantification of NV-CoV-2 in Plasma Samples was done using a validated Direct ELISA as described in the analytical method. For all doses, NV-CoV-2 was detected in rat plasma producing an initial increase that peaked between 4-8 hours. The results show that plasma concentrations decreased to below the level of detection between 24 and 48 hours.

### 2. *In vitro* antiviral activity of NV-CoV-2 (**Figure 3**)

Dose-responsive antiviral effects of NV-CoV-2, NV-CoV-2-R and RDV in SBECD were observed as protection of MRC-5 cells infected with CoV 229E virus (A); and LLC-MK2 cells infected with CoV NL63 (B) virus.

**Figure 3:**
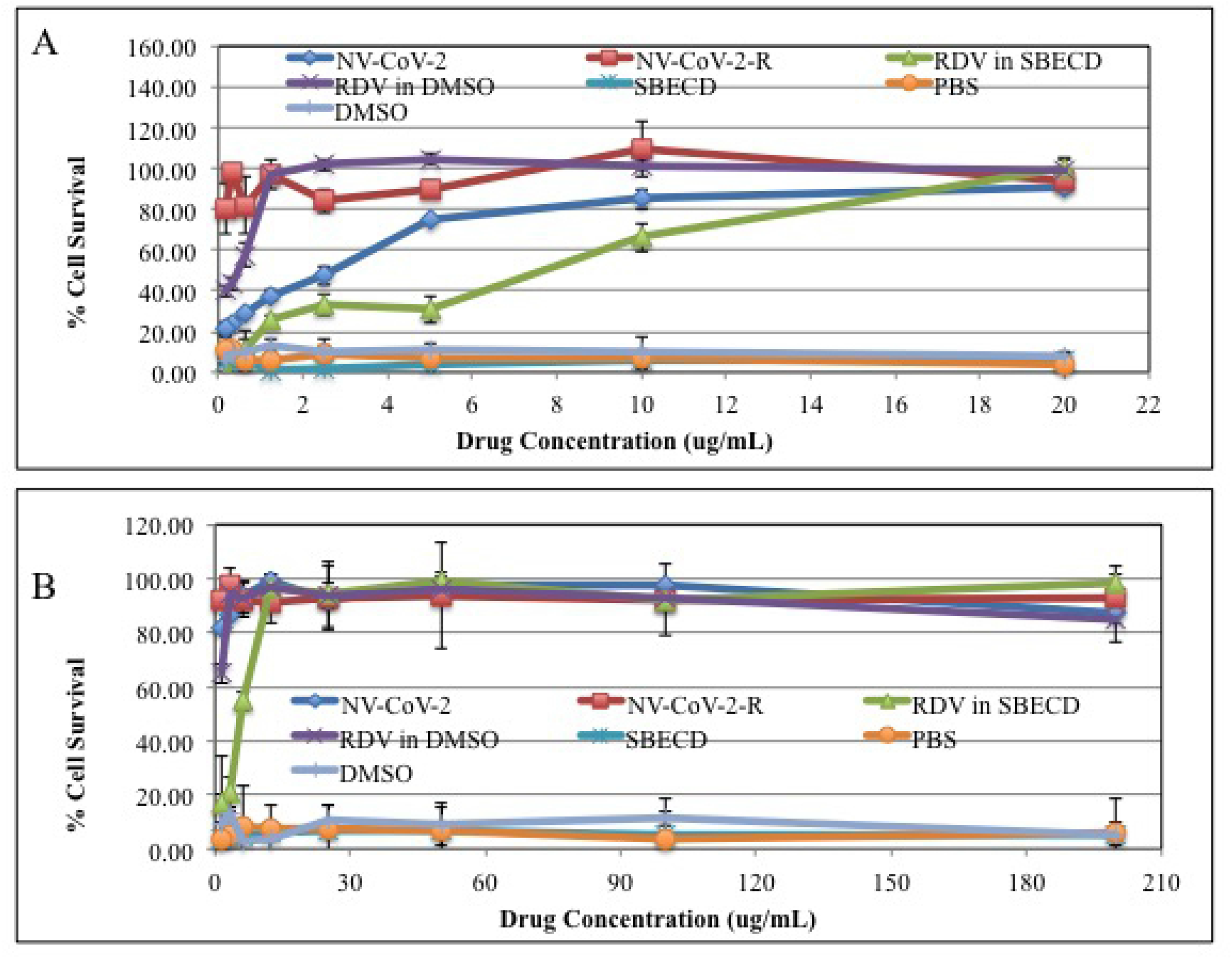
*In vitro* antiviral activity of NV-CoV-2. *In vitro* antiviral activity of the drugs were measured using two cell lines and two different viruses (A: MRC5 cells infected with hCoV-229E; and B: LLC-MK2 cells infected with NL-63 virus) as described in materials and methods. Vehicle control while not showing any antiviral activity, the NV-CoV-2 and NV-CoV-2 are exhibiting their antiviral capacity. As a positive control RDV was used.

### 3. *In vivo* antiviral activity of NV-CoV-2: Days and Dose-response of NV-CoV-2 in male rats infected with CoV-NL63

As shown in the **Figure 4** below, the untreated rats and the vehicle-treated rats infected with the CoV-NL63 virus survived until Day 5. Rats treated daily with RDV only, 10 mg/kg, survived 7.5 days. The rats treated with NV-CoV-2, 160 and 320 mg/kg, survived until Days 13.5 and 14, respectively. Thus, treatment with NV-CoV-2, at both dose levels, markedly extended survival of rats infected intra-tracheally with a lethal dose of human CoV-NL63 virus. Importantly, both treatments were also markedly better than RDV treatment alone.

**Figure 4:**
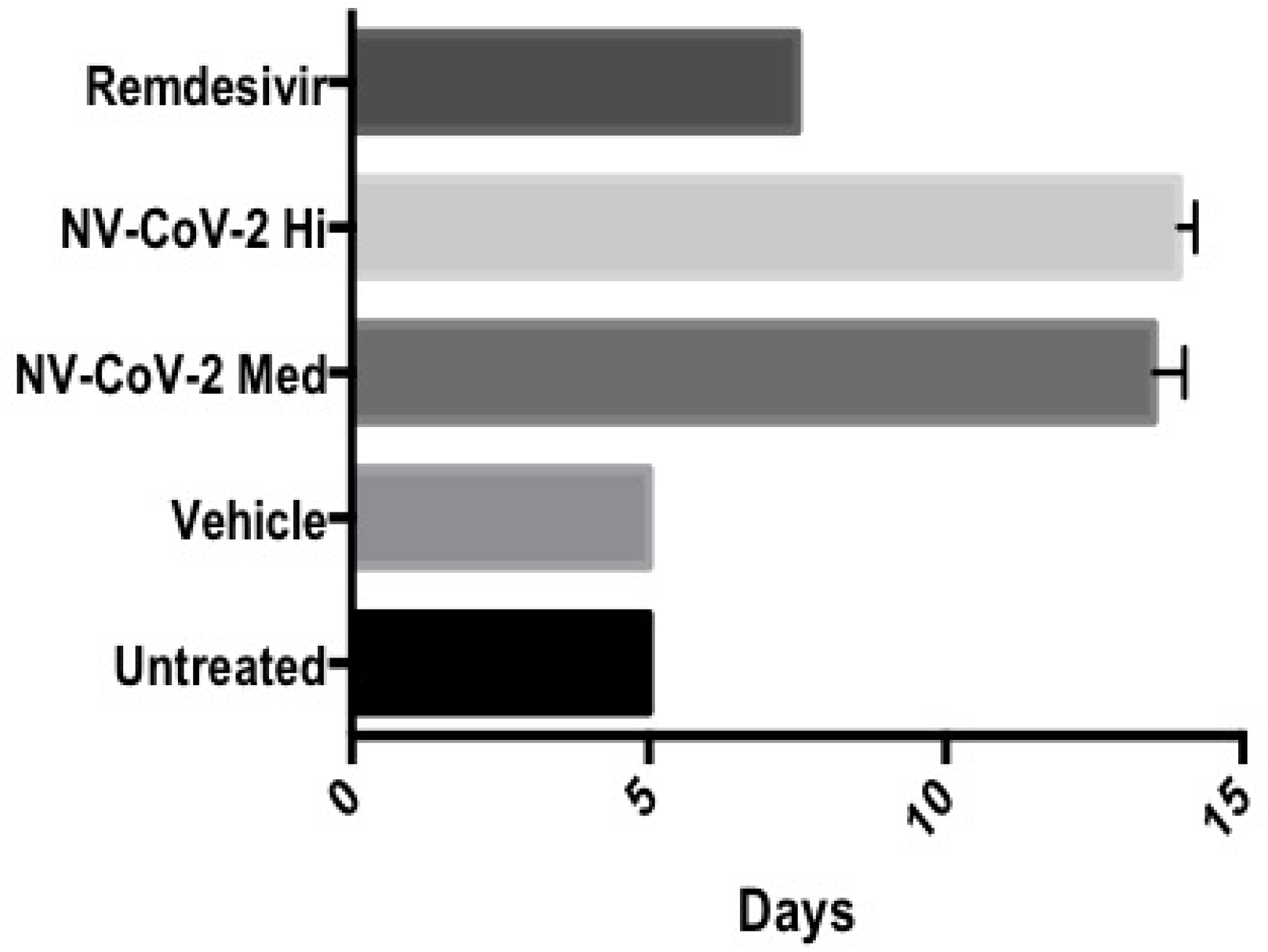
Survival of NV-CoV-2 (Polymer) treated or untreated Rats infected with NL63 virus The untreated rats and the Vehicle-treated rats infected with the Cov-NL63 virus survived until Day 5. Rats treated daily with Remdesivir only, 10 mg/kg, survived 7.5 days. The rats treated with NV-CoV-2, 160 and 320 mg/kg, survived until Days 13.5 and 14, respectively (4A). Further the survival of the infected rats were dependent on the number of days and dosage of the drug, NV-CoV-2 polymer (4B).

Further, the survival of rats following a lethal dose of Cov-NL63 was dependent on the number of days and dosage of the drug used for the treatment **(Figure 5)**. An increase in the number of days of treatment increased the survival days. The survival of rats administered NV-CoV-2 for 1 day was similar to that of untreated rats and administration of vehicle-treated rats, 6- 7 days survival; injection for 3 days and 5 days increased survival to 13.4 days and 17.6 days, respectively.

**Figure 5:**
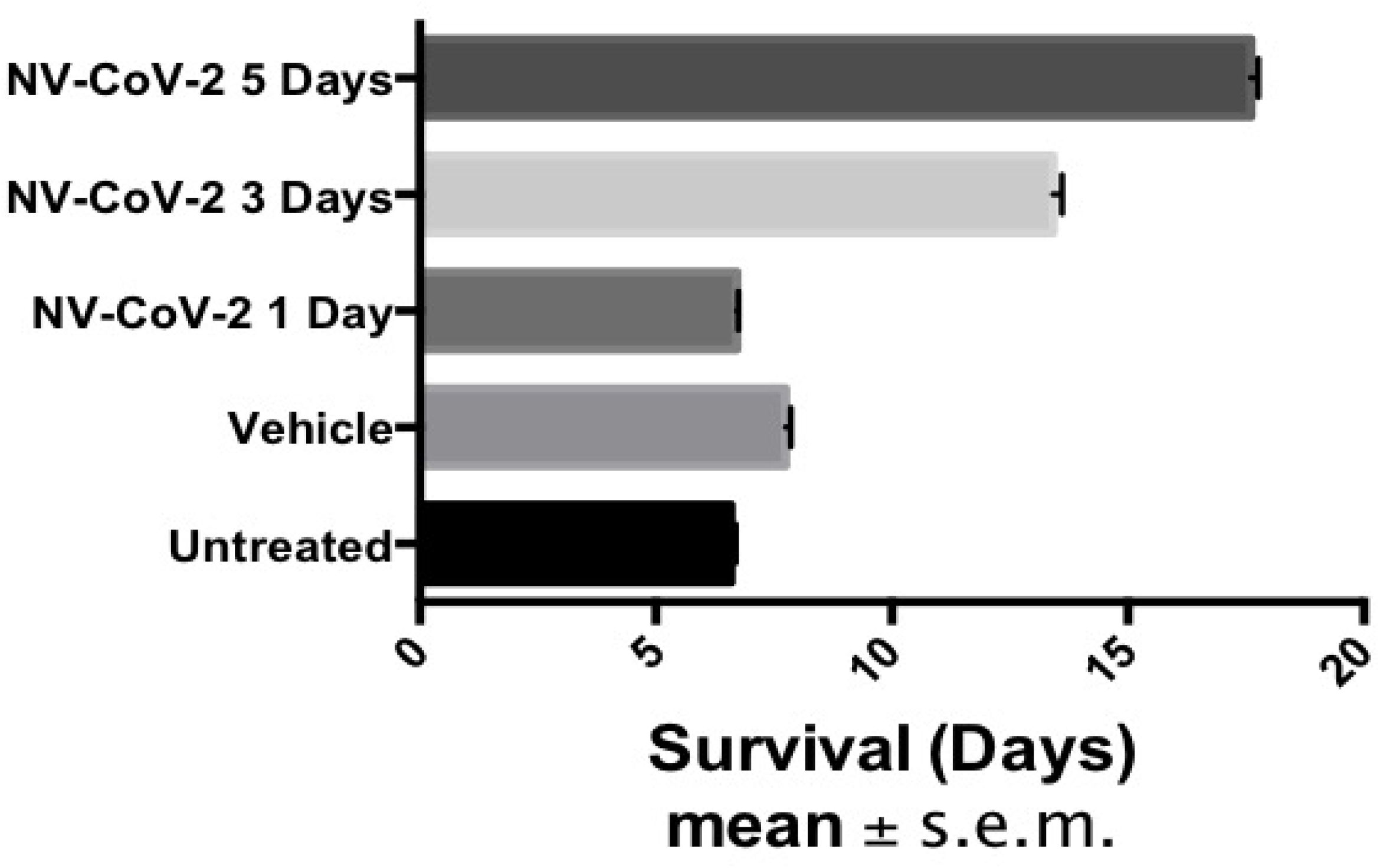
Time dependent antiviral effect of NV-CoV-2 against NL-63 infected animals. The survival rate of NL-63 infected rats were increased as the number of days of treatment was increased.

### 4. The survival of rats after administration of NV-CoV-2-R

for 1 day was slightly increased, 8.8 Days, compared to that of vehicle-treated rats, 7.7 days; injection for 3 days and 5 days increased survival to 13.9 days and 18 days, respectively **(Figure 6)**. Similar to treatment with NV-CoV-2, the increase in survival after 3 days and 5 days of treatment with NV-CoV-2-R was also reflected in the body weight changes over time post-infection.

**Figure 6:**
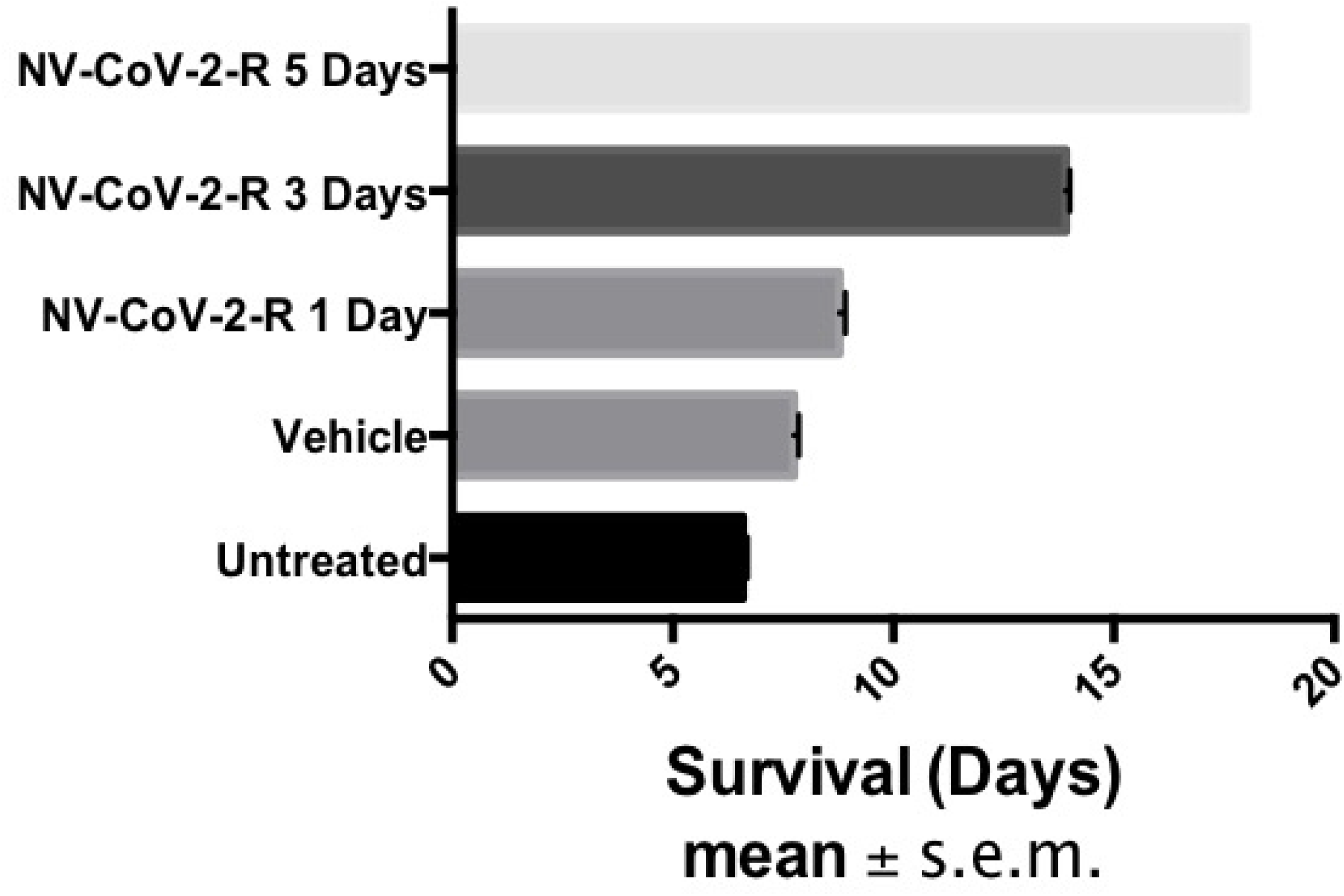
Time dependent. Antiviral effect of Polymer- encapsulated RDV, NV-CoV-2-R against NL-63 infected animals. The survival rate of NL-63 infected rats were increased as the number of days of treatment with the NV-CoV-2-R was increased.

### 5. Body weight measurement

The efficacy of NV-CoV-2, at both dose levels, on survival was also reflected in the body weight changes. Body weight loss increased significantly as the animals became moribund. As shown in the **Table 9** below, by Day 5 post-infection, the infected, untreated rats and the Vehicle-treated rats had a mean body weight loss of 30.8 and 26 g, respectively, while the body weight loss in all other groups ranged between 3.0 and 4.5 g. On Day 7, the daily Remdesivir-treated rats had lost 30.1 g whereas the body weights of the NV-CoV-2- and NV-CoV-2-R-treated groups were not significantly changed from Day 5. By Day 13, both dose groups of the NV-CoV-2-treated rats had lost significant body weight, 25.7 and 12.6 g, respectively.

**Table 9:**
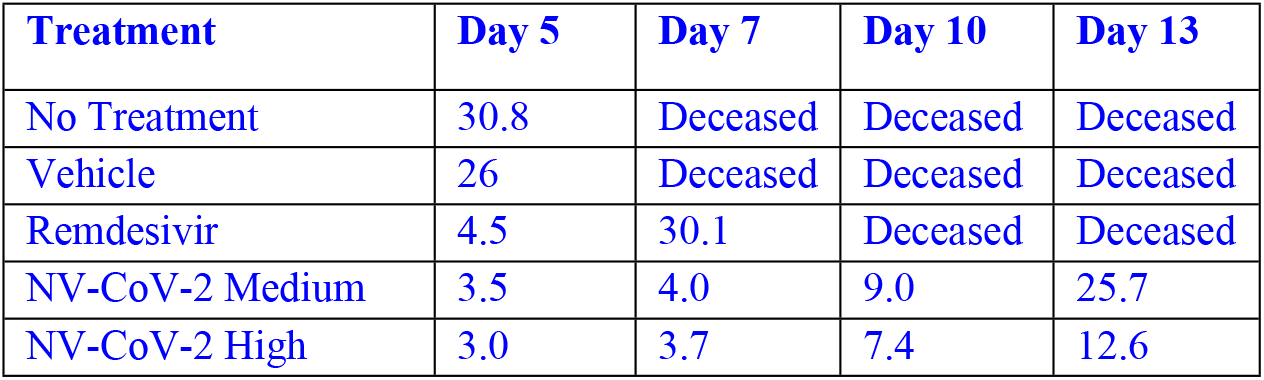
Body Weight Loss (g) Post-Infection.

| **Treatment** | **Day 5** | **Day 7** | **Day 10** | **Day 13** |
| --- | --- | --- | --- | --- |
| No Treatment | 30.8 | Deceased | Deceased | Deceased |
| Vehicle | 26 | Deceased | Deceased | Deceased |
| Remdesivir | 4.5 | 30.1 | Deceased | Deceased |
| NV-CoV-2 Medium | 3.5 | 4.0 | 9.0 | 25.7 |
| NV-CoV-2 High | 3.0 | 3.7 | 7.4 | 12.6 |

Thus, the treatment with NV-CoV-2 at both 160 and 320 mg/kg/injection of Days 0,1,3,5, and 7 had significant effect on survival in rats using a lethal model of infection with the human coronavirus CoV-NL63. Importantly, NV-CoV-2 treatment was clearly superior to daily treatment with *Remdesivir*.

### 6. Efficacy of NV-CoV-2 encapsulated Remdesivir, NV-CoV-2-R, in protecting RDV *in vitro* and *in vivo* from plasma-mediated metabolism

NV-CoV-2 is capable of encapsulating other drugs [4, 5]. This ability was exploited by creating another drug candidate, NV-CoV-2-R, which encapsulates inside NV-CoV-2 the only approved antiviral drug against SARS-CoV-2, RDV [8].

*In vitro,* NV-CoV-2 polymer encapsulation protects RDV from plasma-mediated catabolism **(Figure 7).** *In vivo* study with male rats animal model, the results were shown in **Figure 8** and **Figure 9**. After the 1^st^ injection of NV-CoV-2-R-Med, the accumulation of RDV is greater which is more evident after 5^th^ injections. No such differences were found in the accumulation of RDV from 376-R-SBECD (Gilead) injections, both 1^st^ and 5^th^.

**Figure 7:**
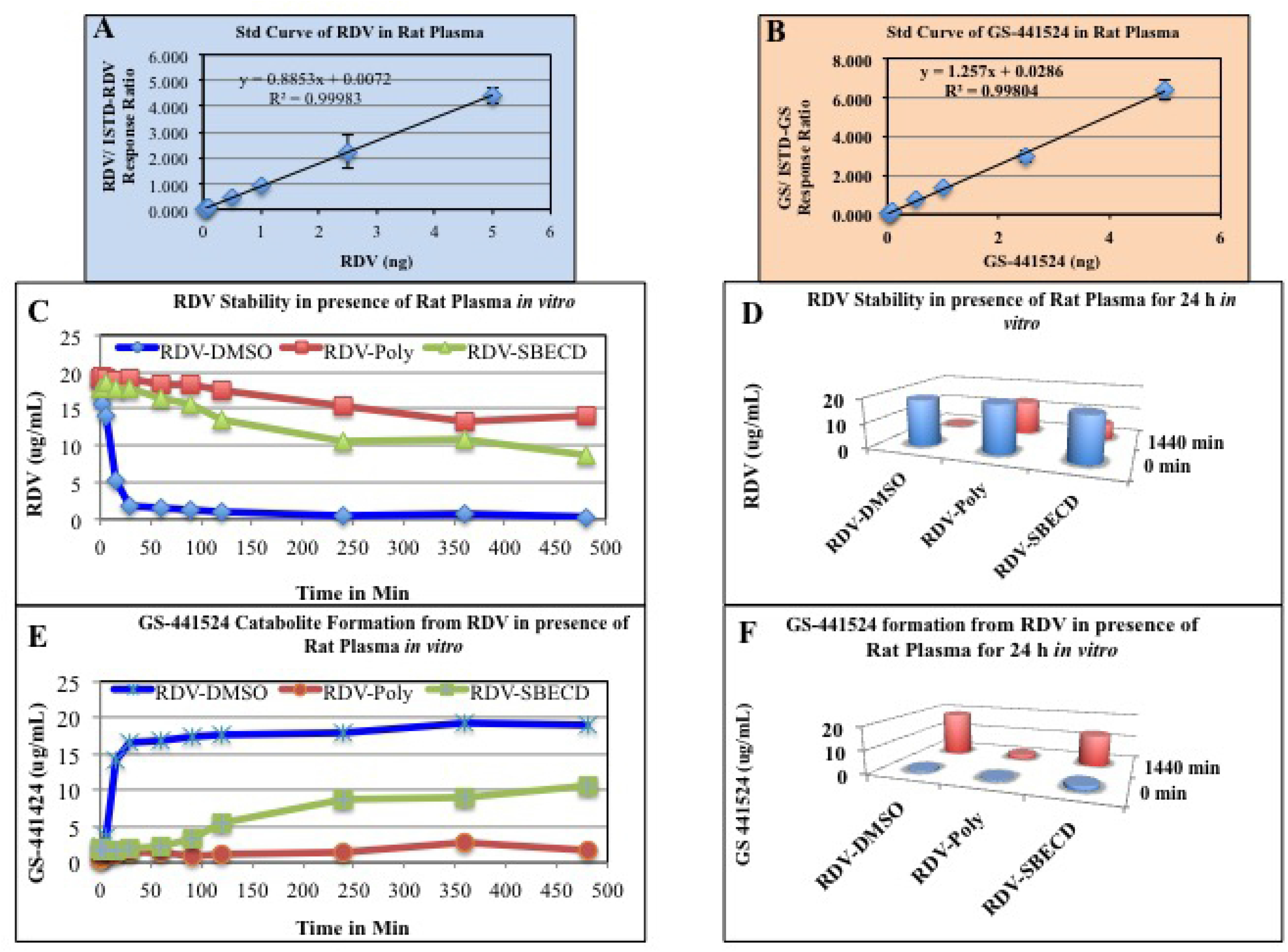
Panels A and B display the Standard Curve of Remdesivir and GS-441524 respectively. Panels C, show the RDV stability in presence of Rat Plasma *in vitro*. The drugs use in this experiments are RDV, Polymer encapsulated RDV (NV-CoV-2-R) and RDV-SBECD (Gilead). Panel D show the values after 24h of the incubation of RDV with the drugs *in vitro*. Panels E, F were showing the values of RDV metabolite, GS441524 as a proof of RDV stability or breakdown *in vitro*.

**Figure 8:**
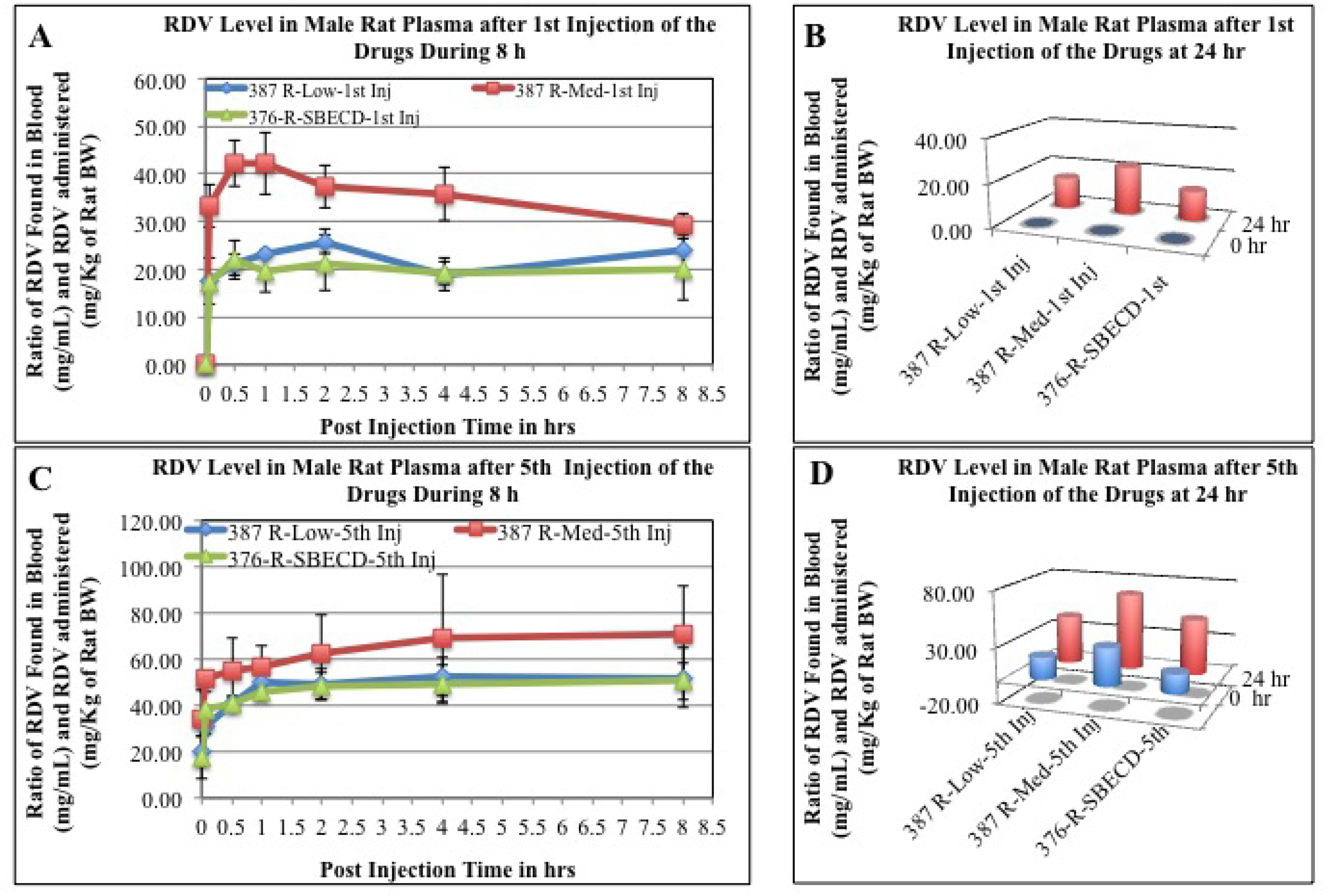
RDV values in Rat plasma after 1^st^ (panels, A, B) and 5^th^ (panels C, D) injection of the drugs. The blood samples collected at different time points after the drug adninistration *i.v.* to the animals, were measured for RDV level as described in the method section. The values obtained as mg/mL were normalized by dividing with the RDV amount administered (mg/kg of rat body weight). Each data point is the mean (±SD) of 3 values and the experiment was repeated three times with similar results.

**Figure 9:**
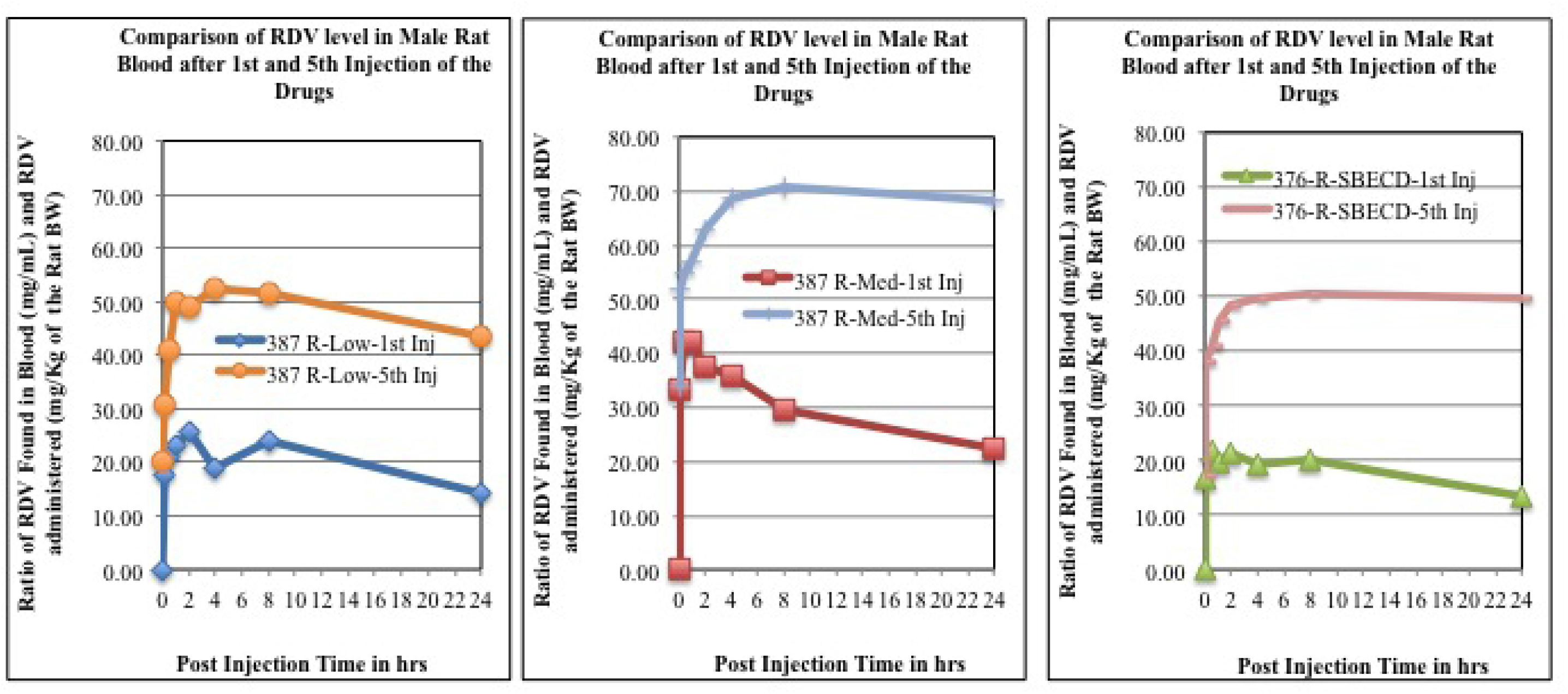
A comparison of RDV level after 1^st^ injection and 5^th^ injections od the drugs were shown.

In brief, NV-CoV-2 polymer can encapsulate *Remdesivir* efficiently. NV-CoV-2-R shows an RDV peak in LC-MS chromatography but the polymer NV-CoV-2 alone does not show (data not shown). Additionally, we tried to measure RDV levels in the respective vehicles, such as DMSO, NV-CoV-2, and PBS. The results are below the detection level (data not shown).

### 7. Respiratory Parameter Results

Effects of the vehicle and NV-CoV-2 (25, 50 and 100 mg/kg) on pulmonary parameters are shown in the **Table 10** below.

**Table 10:** Effects of the vehicle and NV-CoV-2 on pulmonary parameters.

| Intravenous Treatment | Time (hr) | Respiratory Rate (breaths/min) |  | Tidal Volume (mL) |  | Minute Volume (mL/min) |  |
| --- | --- | --- | --- | --- | --- | --- | --- |
|  |  | Mean | SEM | Mean | SEM | Mean | SEM |
| Vehicle Control (mL/kg) | 0 | 133.63 | 9.11 | 1.34 | 0.08 | 174.91 | 12.81 |
|  | 0.25 | 137.72 | 7.99 | 1.53 | 0.10 | 203.24 | 9.20 |
|  | 1 | 142.68 | 12.27 | 1.46 | 0.14 | 197.87 | 9.24 |
|  | 2 | 142.84 | 13.18 | 1.36 | 0.11 | 185.35 | 8.79 |
|  | 4 | 137.05 | 6.01 | 1.51 | 0.05 | 204.08 | 11.29 |
| NV-CoV-2 (25 mg/kg) | 0 | 142.09 | 10.23 | 1.45 | 0.09 | 199.60 | 9.67 |
|  | 0.25 | 142.54 | 16.48 | 1.43 | 0.09 | 199.98 | 20.42 |
|  | 1 | 150.46 | 9.23 | 1.27 | 0.06 | 185.69 | 10.27 |
|  | 2 | 146.38 | 10.62 | 1.29 | 0.07 | 183.53 | 12.49 |
|  | 4 | 139.34 | 7.66 | 1.34 | 0.07 | 183.37 | 10.95 |
| NV-CoV-2 (50 mg/kg) | 0 | 157.58 | 6.35 | 1.33 | 0.12 | 208.67 | 15.97 |
|  | 0.25 | 151.88 | 5.56 | 1.43 | 0.08 | 211.23 | 11.82 |
|  | 1 | 146.85 | 4.29 | 1.37 | 0.10 | 199.68 | 13.94 |
|  | 2 | 146.07 | 8.10 | 1.40 | 0.10 | 201.93 | 20.20 |
|  | 4 | 137.89 | 2.95 | 1.45 | 0.08 | 195.59 | 13.31 |
| NV-CoV-2 (100 mg/kg) | 0 | 126.70 | 10.66 | 1.45 | 0.14 | 189.56 | 26.62 |
|  | 0.25 | 153.77 | 7.01 | 1.54 | 0.08 | 229.26 | 13.39 |
|  | 1 | 152.22 | 10.75 | 1.50 | 0.11 | 220.26 | 11.19 |
|  | 2 | 143.41 | 9.77 | 1.49 | 0.10 | 207.34 | 12.25 |
|  | 4 | 137.21 | 10.93 | 1.45 | 0.08 | 195.24 | 19.10 |

Mean and standard error of the mean (SEM) calculated using non-truncated values. Time zero (0) = pre-dose value. There were no statistically significant ( $p \leq 0.05$ ) changes compared to the vehicle control group.

The intravenous administration of NV-CoV-2 at 25, 50 or 100 mg/kg did not induce any significant effect on respiratory rate, tidal volume or minute volume when compared to the vehicle control group.

### 8. Neurobehavioral Parameter Results

As shown in the **Table 11** below, no apparent neurobehavioral or clinical signs were observed at 1 or 4 hours post-dose in any rats receiving the vehicle. No significant neurobehavioral effects were observed in rats administered 25, 50 or 100 mg/kg doses at the 1- hour post-dose observation. Decreased activity (2/6 animals) and decreased abdominal tone (1-2 animals out of 6) were observed in a small number of animals treated with NV-CoV-2 at 4-hour post-dose observation.

**Table 11:** Neurobehavioral Parameters and Body temperature.

| Group No. | Treatment | Dose (mg/kg) | Volume (mL/kg) | Conc (mg/mL) | Signs Observed (15 Minutes-4 Hours post-dose<sup>a</sup>) | Rectal Body Temperature<sup>b</sup> (°C) |  |
| --- | --- | --- | --- | --- | --- | --- | --- |
|  |  |  |  |  |  | 1 Hr post dose | 4 Hr post-dose |
| 1 | Vehicle control | 0 | 5 | 0 | No signs | 37.9 ± 0.30 | 38.1 ± 0.17 |
| 2 | NV-CoV-2 | 25 | 5 | 5 | 15 Min pd – No signs<br>1 Hr pd - No signs<br>4 Hr pd – Decreased activity (2/6), decreased abdominal tone (1/6) | 38.4 ± 0.15 | 37.6 ± 0.43 |
| 3 | NV-CoV-2 | 50 | 5 | 10 | 15 Min pd – No signs<br>1 Hr pd - No signs<br>4 Hr pd – Decreased activity (2/6), decreased abdominal tone (1/6) | 38.3 ± 0.16 | 38.0 ± 0.25 |
| 4 | NV-CoV-2 | 100 | 5 | 20 | 15 Min pd – No signs<br>1 Hr pd - No signs<br>4 Hr pd – Decreased activity (2/6), decreased abdominal tone (2/6) | 38.8* ± 0.14 | 38.5 ± 0.16 |

<sup>a</sup> Observed for signs of pharmacological or toxicological activity at 15 minutes and 1 and 4 hours after dosing.
<sup>b</sup> Data are presented as mean ± standard error of the mean
\* Statistically significant ( $p \leq 0.05$ ) change compared to the vehicle group – ANOVA followed by a Turkey HSD Multiple Comparison Test (Systat v.9.01)

### 9. Body temperature

was not affected by the intravenous administration of NV-CoV-2 at 25 or 50 mg/kg at 1 or 4 hours post-dose when compared to the vehicle control group **(Table 11)**. Body temperature, 1 hour after the administration of 100 mg/kg dose of NV-CoV-2 was slightly higher (p≤0.05) compared to the control group. Since the difference was marginal (<1.0°C), it may not be considered biologically relevant. Body temperature was not statistically different at 4 hours post-dose by the 100 mg/kg dose of NV-CoV-2.

### 10. Cardiovascular Parameter Results

**Blood Pressure and Heart Rate:** Random occurrence of fluctuations in blood pressure and heart rate are inherent in conscious freely moving animals. Hence consistent (2 or more time points) changes of 15% or more from vehicle controls were considered a significant (biologically relevant) change for blood pressure and heart rate.

No significant effects on blood pressure (systolic, diastolic and mean) or heart rate were observed after the intravenous administration of 25, 37.5 or 50 mg/kg doses of NV-CoV-2. The Figure 10 below shows the mean heart rate and the mean arterial pressure after the 50 mg/kg dose.

**Figure 10:**
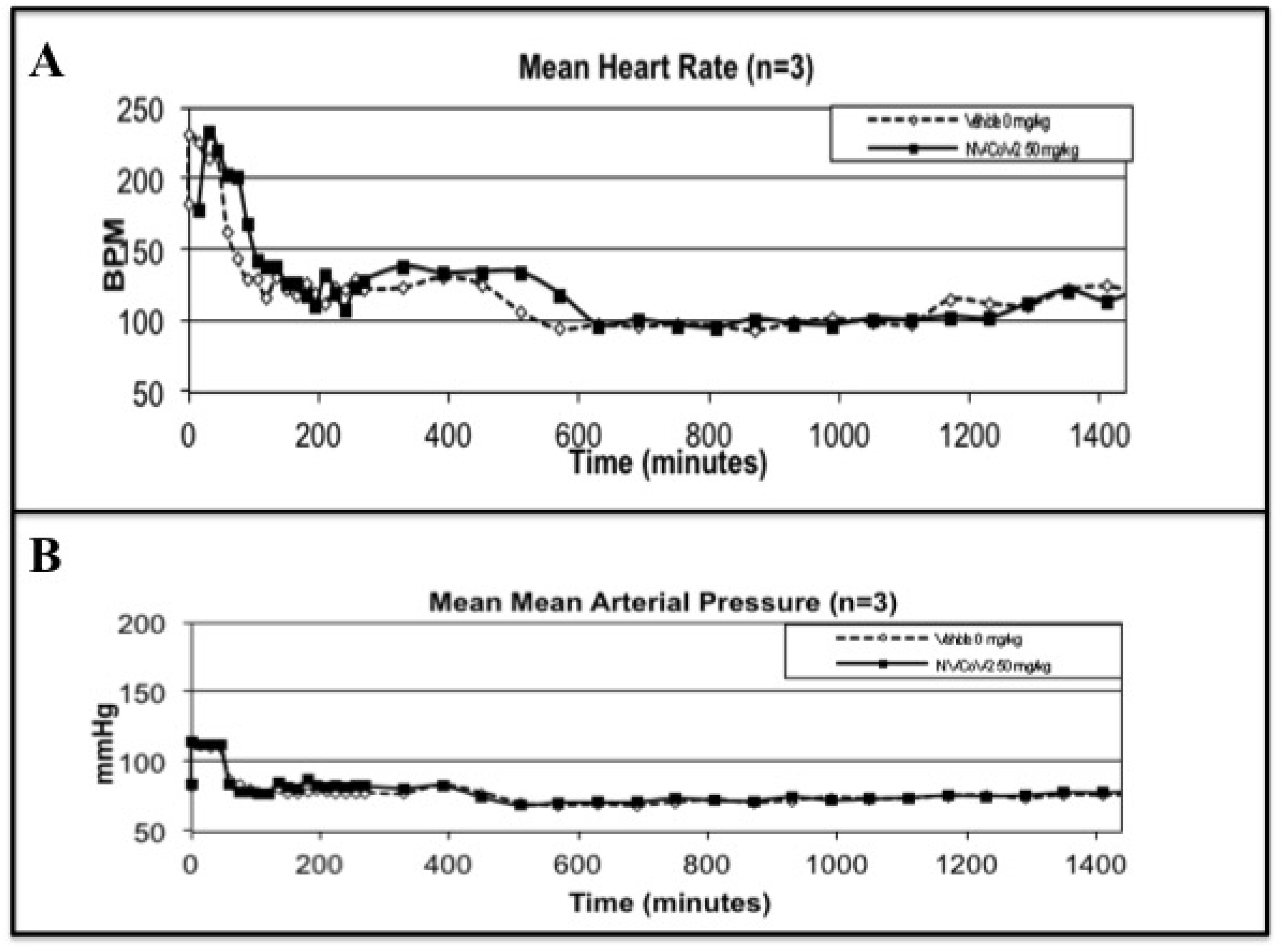
Mean heart rate and arterial pressure after the drug treatment (50 mg) per kg of the male rat body weight.

### ECG

All monkeys maintained sinus rhythm throughout the study. The intravenous administration of 25, 37.5 and 50 mg/kg NV-CoV-2 did not induce any significant effects on quantitative ECG parameters, as measured using the analysis software, in conscious telemetered cynomolgus monkeys.

### Safety Pharmacology Conclusion

The intravenous administration of NV-CoV-2 at doses of 25, 50 and 100 mg/kg did not affect respiratory function in rats. No significant neurobehavioral effects were observed in all dose groups of rats at the 1-hour post-dose observation. Decreased activity (2/6 animals) and decreased abdominal tone (1-2 animals out of 6) were observed in a small number of animals treated with NV-CoV-2 at 4-hour post-dose observation that was not dose-dependent. Body temperature in rats was not affected by the intravenous administration of NV-CoV-2 at 25, 50 and 100 mg/kg.

The intravenous administration of NV-CoV-2 at doses of 25, 37.5 and 50 mg/kg in conscious telemetered cynomolgus monkeys did not induce any significant, biologically relevant effects on heart rate, arterial blood pressure, cardiac rhythm, ECG parameters. All monkeys maintained sinus rhythm throughout the study.

### Immunogenicity

This assay detects all relevant Immunoglobulin (Ig) Isotypes. Since a non-mucosal route of administration was employed and there was no indication of anaphylaxis, the relevant isotypes would be IgG and IgM. The above screening assay is validated to determine lower limit of detection level of isotype IgG and IgM as a group, and is scientifically justified for quantification

of IgG and IgM as a group, using a linear calibration curve. This ELISA has a reliable lower limit of detection of 15ng/ml at which all relevant Ig Isotypes are clearly distinguishable from negative control, and quantifiable with acceptable accuracy and precision.

Immunogenicity determination was done on all animals (4/sex/group) in duplicate following I.V. dosing of NV-CoV-2 in rats with serum samples collected at days 14 and 28 days following the start of the 6 sequential doses over 10 day period. The mean OD of all dose groups is not statistically distinguishable from the vehicle group, both sexes in both Day 14 and Day 28 animals. Under the conditions of the assay method, IgG and IgM antibodies against NV-CoV-2 were not detected and, therefore NV-CoV-2 was not shown to be immunogenic.

## Discussion

NV-CoV-2 is a nanomedicine that we call a *“nanoviricide®”* that is designed to attack most strains of coronaviruses in a broad-spectrum manner. It is composed of a number of antiviral small chemical ligands covalently attached to a micelle-forming polymeric chain. The polymeric chain is a homopolymer from a monomer that is composed of covalently attached polyethylene glycol (PEG), a connecting group, and alkyl pendants. This allows self-assembly of the polymeric chain in aqueous solutions with a PEG shell, a hydrophobic alkyl core, and the ligands being displayed on the outside. The ligand in this case is designed using molecular modeling to bind to the S-protein of SARS-CoV-2 at the same site as where the cognate cellular receptor, ACE2, binds. Thus the *nanoviricide* NV-CoV-2 was designed to neutralize ACE2- binding coronaviruses such as SARS-CoV-1, SARS-CoV-2, and NL63. In practice, it also has activity against human coronavirus 229E (hCoV-229E), which uses Aminopeptidase N (APN) as its cellular receptor, and not ACE2. This appears to be because of certain conserved elements in the quaternary structures of ACE2 and APN that are both peptidases with different specificities. The broad-spectrum activity of NV-CoV-2 against h-CoV-NL63 and hCoV-229E together indicate that it may continue to be active against a number of coronaviruses in spite of mutations.

The mechanism of action of NV-CoV-2 is not well defined but, as described above, the working hypothesis is to bind to free virion particles at multiple points and encapsulate the virus, disabling its ability to infect cells thus neutralizing the virus. This putative mechanism is orthogonal to the mechanism of action of most other anti-coronavirus agents in development **(Fig. 1).** In addition, it may be possible to achieve a stronger antiviral effect by combining NV-CoV-2 therapy with other anti-coronavirus therapies in the future.

The study exhibited no evidence of any adverse reactions in all groups of animals. All groups including the test article groups and the control groups tolerated the compounds with no clinical or behavioral changes observable, no evidence of any deviation from normality. The tolerability of NV-CoV-2 was the same as that of the vehicle control animals. There were no adverse reactions observed while and during the administration of the compounds or during the study period and at postmortem examination. The body fluids and fecal analysis showed no significant difference between the groups, except within the high dosage groups. Histopathological examination showed no changes in any of the organs examined. Based on the results of this study, no Observable Adverse Effects on tolerability could be reported.

At present *remdiesivir* is the only drug that are approved by FDA to be used for COVID- 19 therapy [8]. The standard Veklury® formulation of *remdesivir* in betadex sulfobutyl ether sodium (SBECD) helps with suspending *remdesivir* in solution, but does not appear to be significantly improved upon the metabolic effects. In contrast, NV-CoV-2-R is an encapsulation approach wherein *remdesivir* would slowly leak out into the bloodstream from the polymeric nano-micelle over time, imparting protection against metabolism and sustained effective levels of the encapsulated drug component over a longer time period.

Safety and tolerability of that anti-coronavirus drug candidates (NV-CoV-2 and NV-CoV-2R) were studied here in a rat model and found to be safe and well tolerated. Body weights remained constant and there were no clinical signs of immune or allergic reactions such as itching, biting, twitching, rough coat, etc. Furthermore, there were no observable changes in any organs including large intestine or colon on post mortem in gross histology. This non-GLP safety/tolerability study was conducted under GLP-like conditions by AR BioSystems, Inc., Odessa, Tampa, FL. Further microscopic histology and blood work analyses are in progress [10, 11].

Nonetheless, our platform technology based NV-CoV-2 encapsulated-RDV drug has a dual effect on coronavirus. Our polymer (NV-CoV-2) posses antiviral activity, and protects RDV more than Gilead-RDV, rendering RDV effectiveness against the virus. Furthermore, potential mutations in the virus are unlikely to enable it to escape these drug candidates, as it is a broad spectrum antiviral regimen.

## Acknowledgement

We acknowledge all our colleagues, Secretaries for their help during the preparation of the manuscript by providing all the relevant information.

## Conflict of Interests

Authors, Anil Diwan and Randall Barton are employed by the company *Nanoviricides, Inc*. The remaining authors declare that the research was conducted in the absence of any commercial or financial relationships that could be construed as a potential conflict of interest.

## Authors’ contribution

All the authors contributed equally to prepare this article, read and approved the final manuscript.

## Fundings

Fundings from Nanoviricide, Inc.

## Ethical Statement

Not applicable

## Abbreviations

CoV: Coronavirus;
HCoV: Human Coronavirus
FDA: Food and Drug Administration
hAPN: human Aminopeptidase N;
HAT: Human Airway Trypsin-like protease;
SARS: Severe Acute Respiratory Syndrome;
MERS: Middle East Respiratory Syndrome;
RBD: Receptor Binding Domain
hACE2: human Angiotensin Converting enzyme 2
IBV: Infectious Bronchitis Virus;
TMPRSSII: Transmembrane Protease
SD: Standard deviation
SEM: Standard Error of the Mean
RNA: Ribonucleic Acid;
hDPP4: human Dipeptidyl Peptidase 4;
NV-387: Nanoviricides-Polymer 387
NV-387-R: Nanoviricides-Polymer 387- Remdesivir conjugate
RPL: Rat Plasma
SBECD: Commercial encapsulating agent SBECD (GILEAD)
SBECD-R: Commercial Remdesivir conjugated with SBECD
ADME: Absorption, Distribution, Metabolism And Excretion

## Notes

### Competing Interest Statement

The authors have declared no competing interest.

## References

1. Liu DX, Liang JQ, Fung TS. Human Coronavirus-229E, -OC43, -NL63, and -HKU1 (Coronaviridae). Encyclopedia of Virology. 2021; 428–440. doi:10.1016/B978-0-12-809633-8.21501-X

2. Sharma A, Tiwari S, Deb MK, Marty JL. Severe acute respiratory syndrome coronavirus- 2 (SARS-CoV-2): a global pandemic and treatment strategies. Int J Antimicrob Agents. 2020; 56(2):106054. doi:10.1016/j.ijantimicag.2020.106054

3. https://www.biospace.com/article/releases/nanoviricides-announces-covid-19-clinical-drug-candidate-nv-cov-2-was-effective-against-sars-cov-2-further-demonstrating-its-broad-spectrum-pan-coronavirus-activity/

4. Barton RW, Tatake JG, and Diwan AR. Nanoviricides: Targeted Anti-Viral Nanomaterials Handbook of Clinical Nanomedicine, Nanoparticles, Imaging, Therapy, and Clinical Applications. Eds. Raj Bawa, Gerald F. Audette, Israel Rubinstein. ISBN 9789814669207. Published February 24, 2016 by Jenny Stanford Publishing, pp:1039- 1046

5. Burton R, Tatake JG, Diwan AR. Nanoviricides- A Novel Approach to Antiviral Therapeutics. Bionanotechnology II. *Ed*. David E. Reisner. CRC Press. Taylor and Francis Group, Boca Raton, FL (www.crcpress.com). 2011, pp:141–154

6. Li W, Sui J, Huang IC, et al. The S proteins of human coronavirus NL63 and severe acute respiratory syndrome coronavirus bind overlapping regions of ACE2. Virology. 2007;367(2):367–374. doi:10.1016/j.virol.2007.04.035

7. Chakraborty A and Diwan A. NL63: A Better Surrogate Virus for studying SARSCoV-2. Integr. Mol Med, 2020 doi: 10.15761/IMM.1000408 Volume 7: 1–9

8. COVID-19 Treatment Guidelines. 2020. https://www.covid19treatmentguidelines.nih.gov/antiviral-therapy/remdesivir/.

9. Chakraborty A, Diwan A, Arora V, Thakur Y, Holkar P, Chiniga V. *Nanoviricides* Platform Technology based NV-387 polymer Protects Remdesivir from Plasma-Mediated Catabolism *in vitro:* Importance of its increased lifetime for *in vivo* action. BioRxiv 2021.10.22.465399; doi: https://doi.org/10.1101/2021.10.22.465399

10. https://www.prnewswire.com/news-releases/significantly-improved-safety-profile-andmetabolism-of-remdesivir-observed-due-to-encapsulation-in-nanoviricides-drug-candidate-enabling-potential-highly-effective-pan-coronavirus-antiviral-drug-301382343.html.

11. https://www.marketwatch.com/press-release/nanoviricides-develops-highly-effective-broad-spectrum-drug-candidates-against-coronaviruses-2020-05-12. Nanoviricides Develops Highly Effective Broad-Spectrum Drug Candidates Against Coronaviruses. Published: May 12, 2020 at 7:15 a.m. ET

